# Characterization of quiescent subpopulations and proliferative compartments in glioblastoma

**DOI:** 10.1101/2025.10.03.680278

**Authors:** Anca B. Mihalas, Kelly Mitchell, Sonali Arora, Samantha A. O’Connor, Anoop P. Patel, Christopher L. Plaisier, Patrick J. Paddison

## Abstract

Glioblastoma (GBM) quiescent (Q) cell populations are hypothesized to contain cancer stem-like cells (CSC) that drive tumor growth, cellular heterogeneity, and recurrence. However, GBM tumors do not neatly resolve into developmental hierarchies and Q stem-like activities are difficult to assess. Here, we evaluated tumor Q subpopulations in patient-derived GBM xenograft tumors using live cell reporters, DNA label retention assays, and single cell genomics. Compared to adult neural stems cells (NSCs), GBM Q populations contain hybrid transcriptional states composed of networks found in both dormant and activated adult NSCs, resulting in constitutive expression of key Q egress transcription factors and their targets (e.g., AP-1 and *CCND1/2*). As a result, even the longest Q-residing cells (∼12 days) in xenograft tumors continuously cycle and fail to enter dormant Q states. We provide evidence and hypothesize that transient Q states in primary tumors arise as part of distinct proliferative compartments rather than deterministic developmental hierarchies driven by CSC activity. We further speculate that increases in basal translation rates drive Q instability in GBM tumors.

## Introduction

The concept of cellular quiescence (Q) - a reversible exit from the cell cycle (G0) - has long been recognized as a defining feature of mammalian cell biology. Early studies showed that long-lived adult stem cells, including hematopoietic and neural stem cells (NSCs), can remain dormant for extended periods yet rapidly re-enter the cycle when required (Coller 2019; Cheung and Rando 2013). These observations established Q as an actively regulated state rather than a mere absence of proliferation. Transcriptional, epigenetic, metabolic, and signaling circuits jointly govern entry, maintenance, and exit from Q, preserving stem-cell integrity and enabling tissue regeneration (Urbán et al. 2019; Rodgers et al. 2014).

Single-cell genomic analyses have since mapped Q-to-activation (A) trajectories across adult stem-cell systems. In the neural lineage, Llorens-Bobadilla et al. (2015) used scRNA-seq to show that subventricular-zone NSCs span a continuum from dormant Q to A states: dormant Q NSCs upregulate stress-resilience and metabolic suppression programs, whereas A NSCs induce growth-factor signaling, translation, and cell-cycle drivers (Llorens-Bobadilla et al. 2015). Subsequent single-cell studies corroborated these state transitions in adult NSCs (Borrett et al. 2020; Shin et al. 2015; Dulken et al. 2017). Analogous Q-to-A continua have been documented in hematopoietic stem cells (Wilson et al. 2008; Cabezas-Wallscheid et al. 2017), muscle satellite cells (Machado et al. 2017; van Velthoven and Rando 2019), and intestinal stem cells (Haber et al. 2017; Grün et al. 2016). Collectively, these studies underscore Q as a conserved, dynamic feature of adult stem-cell biology, which limits replication-associated DNA damage, buffers oxidative stress, and reduces cumulative mutational burden (Cheung and Rando 2013; Li and Clevers 2010).

In oncology, analogous Q or slow-cycling tumor populations have been implicated in therapy resistance. In glioblastoma (GBM), the most common and aggressive primary brain tumor in adults, recurrence after standard-of-care chemoradiation is nearly universal, and a long-standing hypothesis posits that a subset of glioblastoma stem-like cells (GSCs) can enter Q states that are refractory to DNA-damaging agents and radiation before re-engaging proliferation to drive relapse (Bao et al. 2006; Lathia et al. 2015).

Evidence supporting this view has accumulated. Q/slow-cycling GSCs are enriched in vivo and in patient-derived models after temozolomide and radiation, consistent with selective survival under cytotoxic stress (Gimple et al. 2022; Deleyrolle et al. 2011). Transcriptomic and functional assays indicate that these Q subpopulations resist genotoxic therapies yet retain tumor-initiating capacity upon reactivation (Suva et al. 2014; Patel et al. 2014). In vivo lineage-barcoding further suggests that GBM tumors can be subdivided into slower- and faster-dividing populations, with the former showing more stem-like transcriptional features (Lan et al. 2017). How Q states intersect with these proliferative compartments to shape tumor heterogeneity and dynamics remains unresolved, and precise delineation of Q populations continues to be a technical challenge.

To address these questions and assess Q states in GBM tumors, we combined GSC xenograft models with live-cell Q reporters together with computational approaches for defining Q-like states in GBM in single cell genomic datasets. Our results indicate that GBM Q is a transient, hybrid state rather than a reservoir of dormant stem cells. We propose that proliferative heterogeneity organized into dynamic compartments, not fixed CSC hierarchies, likely underlies GBM tumor cell diversity.

## Results

### Single cell RNA-seq analysis of live Q populations in X-GBM tumors

To functionally define GBM tumor cells in short- and long-term Q states, we used two orthotopic brain tumor models derived from human patient-derived GBM stem-like cells (GSCs)(GSC-0827 and GSC-0464T)(Mihalas et al. 2025) in conjunction with scRNA-seq and a reporter for Q (p27-mVenus) and also label retention (Dox-H2B-mCherry).

We and others have used p27-mVenus proteolytic reporter for assessing Q states in GSC, NSCs and other mammalian cell types (Oki et al. 2014; O’Connor et al. 2021; Mihalas et al. 2025). In S/G2/M phases of the cell cycle, p27 is targeted for proteolysis by the SCF^Skp2^ E3 ubiquitin ligase complex, while in G1 it is additionally targeted for degradation by the Kip1 ubiquitylation-promoting complex (Coats et al. 1996; Chu et al. 2008). As a result, p27 only accumulates during G0, where it inhibits the activity of Cyclin D–CDK4/6 and Cyclin E–CDK2 complexes (Polyak et al. 1994; Toyoshima and Hunter 1994) . To ensure that the p27 reporter does not have inhibitory activity a p27 allele was used that harbors two amino acid substitutions (F62A and F64A) that block binding to Cyclin/CDK complexes but do not interfere with its cell cycle-dependent proteolysis (Oki et al. 2014; Mihalas et al. 2025). As a result, the p27-mVenus^hi^ population will be composed of cells entering Q-states (e.g., just after mitosis) and those already in short- or long-term quiescence.

We also employed “label retention” assays in which DNA is labeled with H2B-mCherry using a doxycycline inducible system to enable “chase” without dox (Furutachi et al. 2015; Tumbar et al. 2004). This system reads out division history of post-chase, DNA labeled cells, with label diluted two-fold each division.

In addition we utilized a single cell RNA-seq cell cycle classifier (ccAF), which was trained on human NSC scRNA-seq data (O’Connor et al. 2021). In contrast to other cell cycle classifiers (e.g., ccSeraut), ccAF can computationally define Q-like states (dubbed Neural G0) utilizing genes expressed in adult Q NSCs and fetal radial glial (RG) cells, along with absence of cell cycle genes.

We used GSC-0827 (female; recurrent GBM; proneural)(Son et al. 2009; Toledo et al. 2015) and GSC-464T (male; primary GBM; proneural)(Joo et al. 2013) GBM isolates to generate xenograft models. For these models, we have previously created scRNA-seq “reference” data sets which we used for comparison here (Mihalas et al. 2025) (Fig. 1A; Supplementary Fig. 2A). Uniform manifold approximation and projections (UMAP) was used for dimensional reduction of data and generation of *de novo* cell-based clusters (Becht et al. 2018). 9 cell clusters were apparent in the GSC-0827 tumor reference (Fig. 1A) and 15 in the GSC-464T reference (Supplemental Fig. S2), which corresponding cell clusters in p27^hi^ tumors cells (Fig. 1B; Supplemental Fig. S2B), along with the relative frequencies of cells within each cluster (Fig. 1E and F).

**Figure 1:**
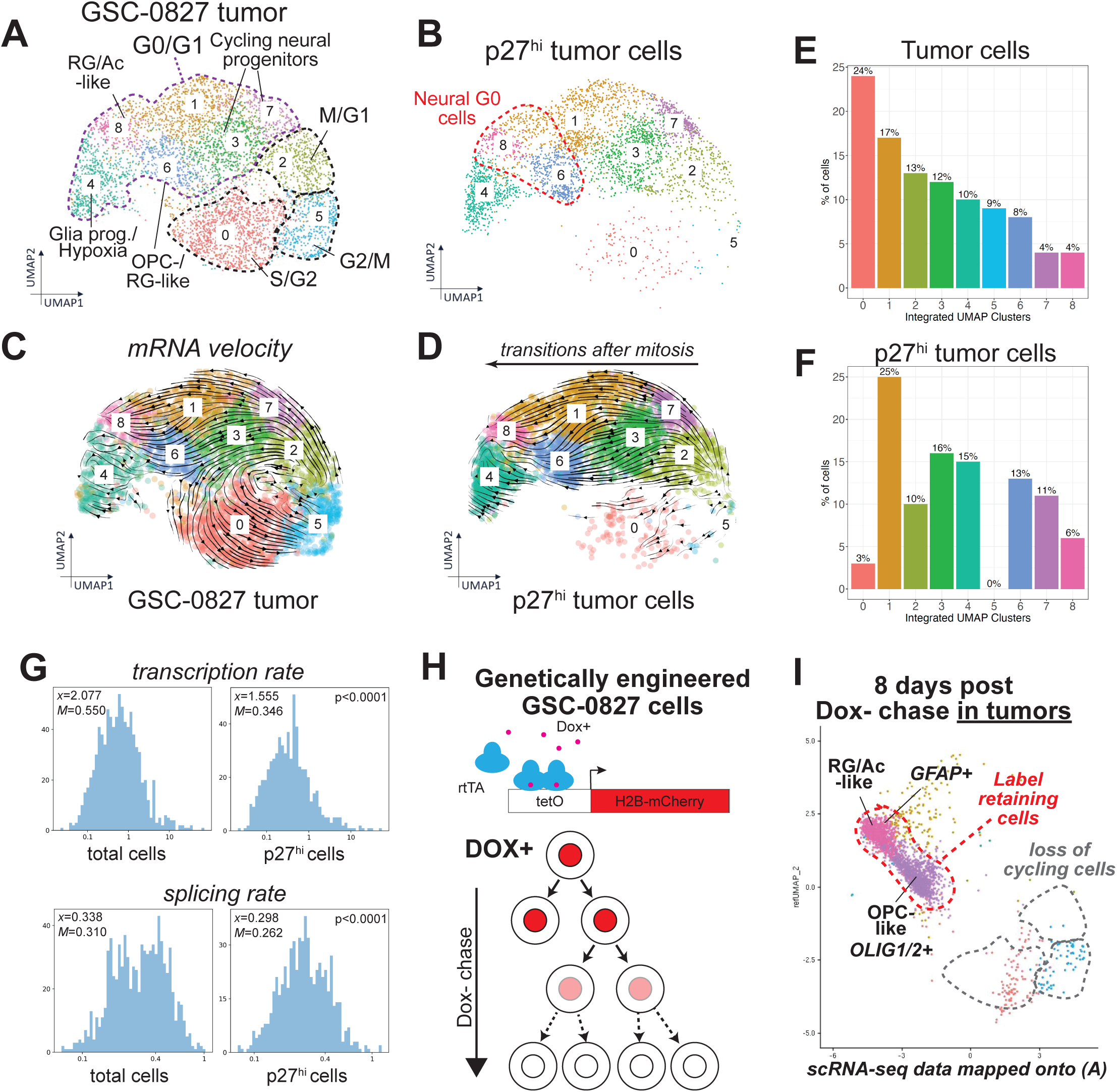
Characterization of GBM Q states using live Q reporters in X-GBM tumors. **A-B.** scRNA-seq UMAPs for GSC-0827 tumor reference and p27hi tumor cells. two human GBM stem-like cells (GSC) tumor models. Overview of experiment, filter cutoffs, and QC analysis are available (Mihalas et al. 2025). **C-D.** mRNA Velocity analysis (scVelo) of p27hi cells reveals trajectories toward Q states after mitotic exit. **E-F.** Relative proportions of mapped single cells appearing in clusters from **A** and **B.** **G.** Histograms of estimated transcription rates and splicing rates for genes for which scVelo dynamic models of transcription and splicing are available (n=1032 genes for total tumor cells; n=749 for p27^hi^ cells)(*x*=mean; *M*= median). A two sample Kolmogorov–Smirnov test was used to test significance. **H-I.** scRNA-seq of X-GBM tumors label retaining cells. The experiments confirm termination points of velocity analysis from A. QC cut offs for scRNA-seq analysis is available in Supplemental Figure S1. Analysis of GSC-464T X-GBM tumors are shown in Supplemental Figure S2. Examples of velocity genes and cell cycle associated TF as shown in Supplemental Figure S3.

To associate clusters with specific cell cycle and developmental states, we performed multiple comparisons of genes differentially expressed within each cluster. This included: differential gene expression and gene set enrichment analysis for each cluster (Mihalas et al. 2025).

Both tumors are highly proliferative with ∼45% of tumor cells predicted in the cell cycle (Fig. 1A; Supplemental Fig. S2) and also have multiple Q/G1 populations that are not apparent in *in vitro* cultures (see (Mihalas et al. 2025)). For GSC-0827 these include OPC- and RG/Astrocyte-like clusters, a hypoxia cluster, and clusters associated with dividing neural progenitors, highlighting that GSCs contain the ‘potential’ to diversify into additional cell states in an orthotopic microenvironment.

The data reveals points of cell cycle ingress and egress for these tumors. For example, for GSC-0827 we could assign cell division phases S/G2, G2/M, and M/G1 to tumor reference clusters (0, 5, and 2, respectively) using the ccAF classifier (Fig. 1A). p27^hi^ tumor cells show dramatic loss of S/G2 and G2/M populations, consistent with loss of actively dividing cells in p27^hi^ cells. However, the M/G1 population remains comparable to the total cell population. This indicates that p27^hi^ populations capture cells exiting the cell cycle and entering transient or long-term quiescent states with low G1 cyclin/CDK activity.

Examination of mRNA velocity lines, which model the direction and speed of cell dynamics based on kinetics of transcription (Bergen et al., 2020), revealed that tumor cells reenter the cell cycle from clusters 2 (M/G1), 3 (cycling progenitors), or 6 (OPC) for GSC-0827 tumors (Figs. 1C and D). However, cluster 1 cells (associated with cycling neural progenitors) also show expression of the G1/S cyclin, CCNE2, DNA replication genes expression (G1/S) (Mihalas et al. 2025), and E2F1 expression (Supplemental Fig. S3). The latter is part of a sequenced expression of cell cycle transcription factors, which includes MYBL2, FOXM1, and MYC, each of which promotes phase-specific cell cycle gene expression, starting with E2F1 (Fischer et al., 2022) (Supplemental Fig. S3).

Consistent with p27-mVenus+ having little or no cyclin/CDK activity, p27^hi^ cells do not reenter the cell cycle based on this analysis (Fig. 1D; Supplemental Fig. S2). Instead, GSC-0827 cells transition from M/G1 along two general tracks, either through cluster 3 to 6 (OPC) and then 8 (RG/Ac) or through clusters 3 or 7 to 1 and then 8. Of note, cells along the 7-1-8 cluster track express CD44 (a marker of mesenchymal GBM cells) and appear largely mutually exclusive from cells that express OLIG1 (an OPC marker) (Mihalas et al. 2025). However, both trajectories terminate in an RG-like cluster. Because we know where cells exit the cell cycle in this analysis, the data support a model whereby tumor cells with longer resident times in Q-like states ultimately adopt OPC-like or RG-like transcriptional features. A similar scVelo trajectory was noted in GSC-464T tumors (Supplemental Figure S2). p27^hi^ populations also in general displayed lower rates of transcription and splicing (Fig. 1G). Lower transcription rates and RNA content are hallmarks of quiescent cells (McKnight et al. 2015; Darzynkiewicz et al. 1980).

To address this point, we performed scRNA-seq analysis of H2B-mCherry label retaining GSC-0827 tumors cells 8 days after Dox chase. At this time point, LR cells are estimated to represent ∼15% of total tumor (see below). Confirming predicted Q populations, these cells map onto the RG/Ac/OPC-like populations with highest proportions of ccAF-predicted Q cells and termination points of scRNA-seq velocity lines (Fig. 1H and I).

### X-GBM Q populations contain hybrid Q and A NSC transcriptional networks

To further, analyze Q populations in X-GBM tumors, we discovered TF regulators of the marker genes from GSC-0827 cluster 8 using TFRegMap. In this analysis, transcription factor activity is inferred by enrichment of TF target genes (Plaisier et al. 2016) and a significant correlation between the TF regulator with the eigengene of the TF targets (first principal component corrected for sign). For GSC-0827 cluster 8, 16 TFs were highly enriched for among top 197 genes expressed in human radial glial from ToppCell single cell gene atlas (FDR=7.489E-17) (Jin et al. 2021). While not unexpected based on our previous analysis, these TFs, however, also included ones that we would not expect to be present in Q cells, including AP-1 and ERG transcription factors.

These factors have well established roles in promoting Q egress and cell cycle entry (Gashler and Sukhatme 1995; Pardee 1989; Wisdom et al. 1999; Molnar et al. 1994). Consistent with this notion network analysis using cancer hallmark categories revealed their associations with self-sufficiency in growth signals, insensitivity to antigrowth signal, evading apoptosis, etc. (Fig. 2A).

**Figure 2:**
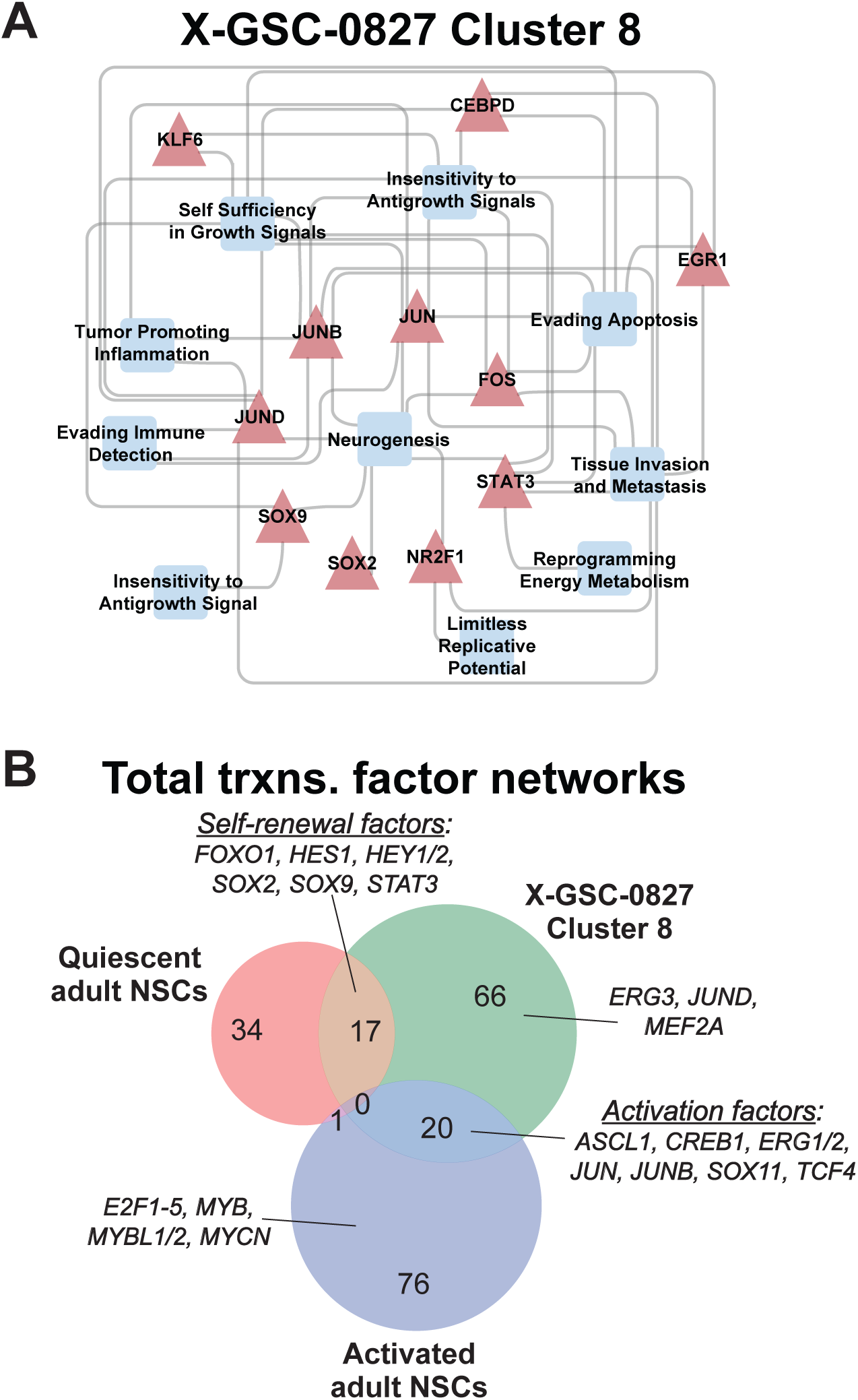
Transcriptional network analysis of X-GBM Q populations. **A.** Network analysis from gene set enrichment analysis cluster 8 from GSC-827 X-GBM for Cancer Hallmarks. scRNA-seq cluster marker genes (log2FC ≥ 0.50; p_val_adj ≤ 0.05) were used to identify enriched TF motifs within cluster 8 (Hypergeometric enrichment; FDR<.05). The eigengene for TF targets used identify TFs with expression that correlates targets (abs(R) ≥ 0.3, FDR corrected p-val ≤ 0.05) (**Supplemental Table S1**). GO term enrichment analysis was then used to identify cancer hallmarks with (p ≤ 0.01) semantic similarity score of ≥ 0.80 (**Supplemental Table S2**). **B.** Overlap of TF network activity in Q and A NSCs from Llorens-Bodadilla et al. (2015) and cluster 8 from GSC-827 X-GBM using TFRegMap analysis (**Supplemental Table S3**).

To further explore these results, we compared them to transcription networks associated with Q and A NSCs. To this end, we next discovered TF regulators of marker genes from the Q and A adult mouse NSCs through analysis of NSC scRNA-seq data sets available from Llorens-Bobadilla et al. (2015) (Fig. 2B). We identified 53 and 97 candidate TF regulators from Q and A NSCs, respectively, which were nearly mutually exclusive. The Q regulatory network was enriched for TFs implicated in NSC maintenance or self-renewal (e.g., FOXO1, HES1, HEY1/2, SOX2, SOX9, and STAT3), while the A regulatory network harbored key cell cycle and growth responsive TFs (e.g., E2F1, ERF1, JUN, MYB, MYCN). Next, we found 17 transcriptional regulators overlapping between the X-GBM 0827 cluster 8 and mouse Q NSC regulatory networks, including the ones called out above. Not surprisingly we also observed an overlap of 20 TF regulators between the X-GBM −827 cluster 8 and mouse A NSC regulatory networks, which include ERG1/2, JUN, and JUNB among others (Fig. 2B).

Because these “activation” TFs are well known for roles in promoting Q egress and cell cycle gene expression, we further investigated cyclin and AP-1 transcription factor expression in NSCs and X-GBM subpopulations from single cell data. It has been well established that cyclins are cell cycle regulated in a pattern shown in Figure 3A (Darzynkiewicz et al. 1996; Sherr 1995). Expression of D type cyclins has been used to discriminate Q from cycling cells which have been mitogenically stimulated to enter the cell cycle (Darzynkiewicz et al. 1996). Cyclin D expression is controlled by a number of transcription factors associated with NSC A networks, including AP-1 and ERG.

**Figure 3:**
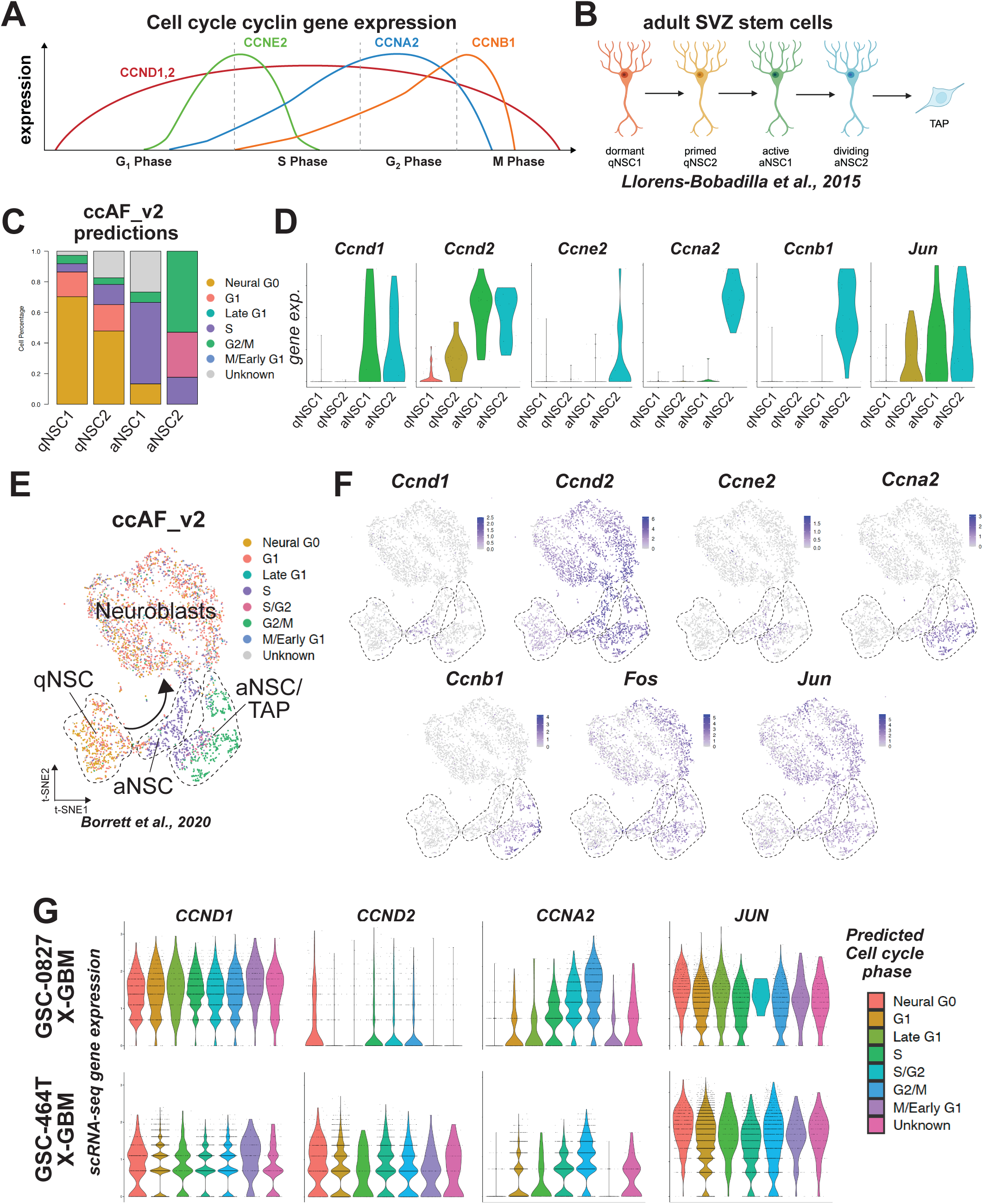
Comparisons of Cyclin and AP-1 gene expression in adult NSC and X-GBM tumors. **A.** Cyclin gene expression across the cell cycle. **B.** Stages of adult SVZ-derived NSC activation as defined by Llorens-Bobadilla et al. (2015). **C.** ccAF_2 cell cycle predictions for qNSCs and aNSCs. **D.** Cyclin and Jun expression among qNSCs and aNSC populations. **E.** UMAP of adult SVZ-derived NSCs from Borrett et al. (2020) classified using ccAF. **F.** Cyclin, Fos, and Jun expression among using UMAP from **E**. **G.** CCND1, CCND2, CCNA2, and JUN expression among ccAF-defined subpopulations of GSC-0827 and GSC-464T xenograft brain tumors. Gene expression is also broken down by de novo UMAP cluster and also shown for p27hi and H2B-mcherry-retaining populations in Figure SX, showing similar results.

We first wished to establish a baseline of expression for Q NSC populations, which should show minimal expression of both Cyclins and activating TFs. To this end we analyzed two adult mouse (subventricular zone) NSCs data sets, adopting their naming conventions for Q and A NSC: qNSC1, qNSC2, aNSC1, and aNSC2 for Llorens-Bobadilla et al. (2015) (Fig 3B, 3C, and 3D) and qNSC, aNSC, and TAP for Borrett et al. (2020) (Fig 3E and 3F). This was indeed the case, Q populations showed little or no expression of Cyclin D (and other cyclins) or AP-1 factors, while A populations showed expression of D and phase-specific cyclins, along with AP-1 factors.

We next examined X-GBM scRNA-seq using ccAF cell cycle phase categories and de novo clusters for tumor reference, p27hi and LR cells. The analysis demonstrated that in predicted and functionally defined Q populations, cyclin D and AP-1 expression remain similarly high to cycling cells.

To further establish the notion that GBM Q states likely exhibit Q+A hybrid NSC TF networks, we analyzed both cyclin D expression and AP-1 expression in scRNA-seq data from 6 primary GBM tumors (Fig. 4A). As with X-GBM tumors, these data reveal that ccAF predicted Q states have constitutive expression of cyclin D and also JUN. TF network predictions from these populations revealed a similar result to X-GBM Q cells, with overlap of Q TF networks and pro-proliferative A TF networks (Fig. 4B). They also harbor predicted pro-proliferative networks not observed in NSCs (that have been previously associated with GBM proliferation (e.g. GLI2 (Huang et al. 2018), HOXA/PBX1 (Arunachalam et al. 2022), LEF1 (Kahlert et al. 2015), ONECUT2 (Haddadi et al. 2024), ZNF217 (Mao et al. 2011), etc.).

**Figure 4.**
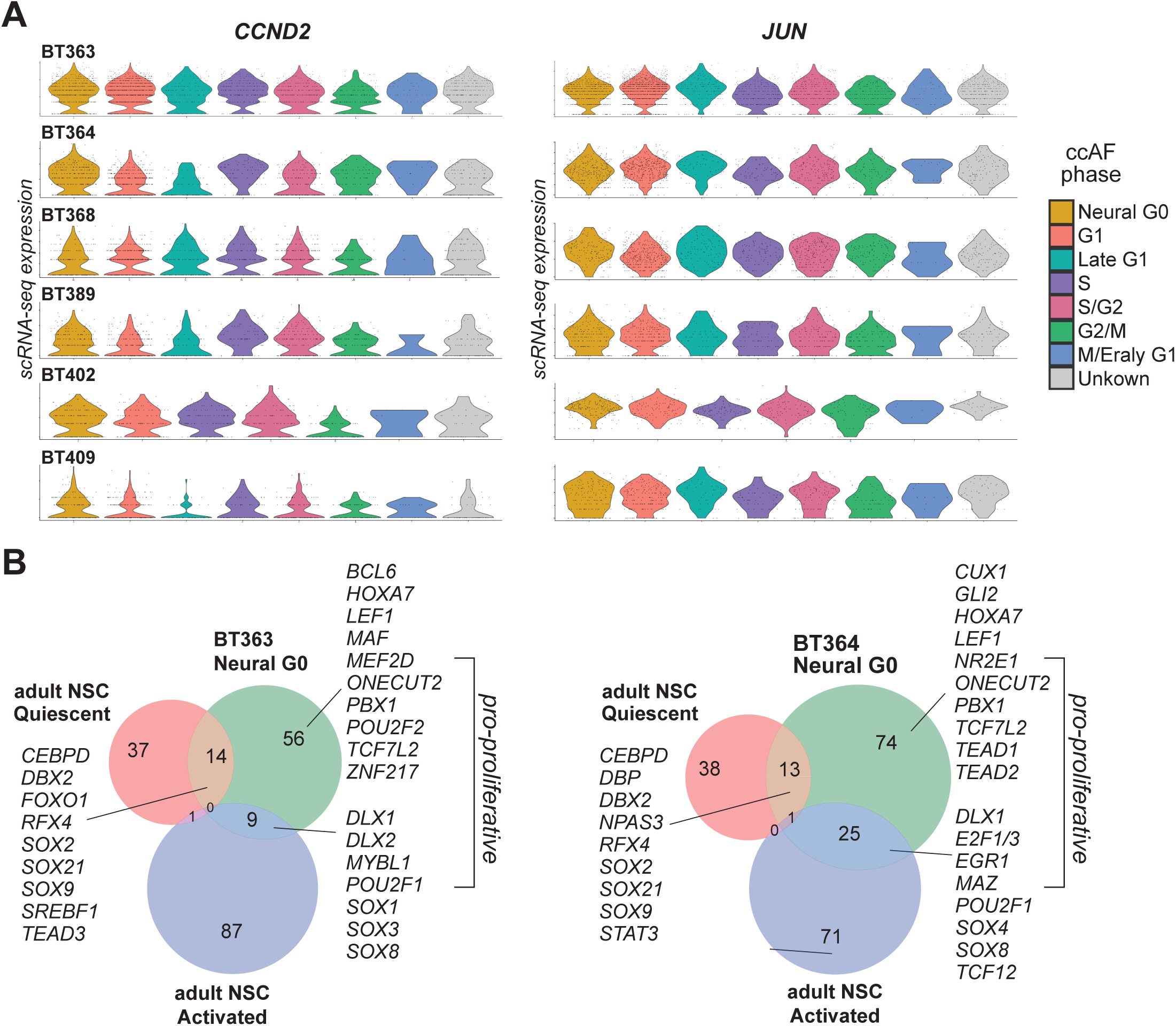
Examination of cell cycle states of among subpopulations of X-GBM tumors. **A.** Expression of CCND2 and JUN among ccAF-defined subpopulations of scRNA-seq data from 6 primary GBM tumors from Couturier et al. (2020). **B.** Overlap of TF network activity in Q and A NSCs from Llorens-Bodadilla et al. (2015) and ccAF-defined Neural G0 cells from BT363 and BT364 tumor samples from **A** using TFRegMap analysis (**Supplemental Table S4**).

This suggests that X-GBM and primary GBM Q cells likely have only transient Q and remain in a hybrid Q-A stated primed for cell cycle reentry.

### X-GBM tumors exhibit continuous dilution of DNA labels

To investigate the phenotypic behavior of candidate Q populations, we performed in vitro and in vivo DNA label retention (LR) assays using H2B-mCherry and EdU pulse-chase methods. In these assays, cells that either enter long-term quiescence (e.g., Q NSCs) or undergo terminal differentiation retain the DNA label after the pulse period (Fig. 5A). If cells fail to do either and continue to proliferate, continuous dilution of label is observed (Fig. 5B).

**Figure 5:**
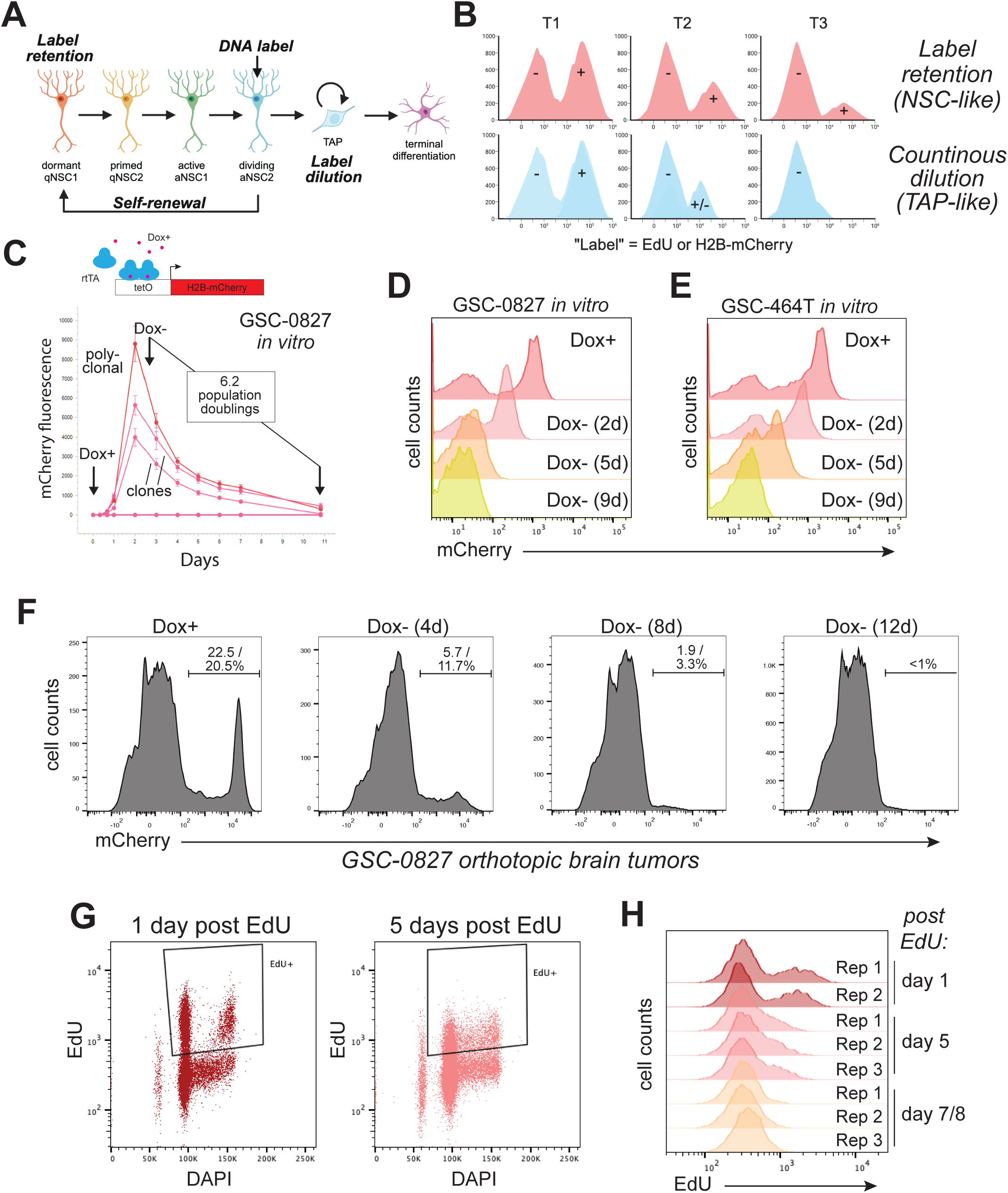
Label retention experiments in X-GBM tumors. **A.** Model for label retention vs. label dilution in qNSCs. **B.** Model FACS data indication label retention vs. label dilution. **C.** In vitro validation studies of label retention using Dox-H2B-mCherry GSC-0827 cells using 2 LV-Dox-H2B-mCherry clones or a FACS sorted poly clonal population. Dox was administered for 48hrs and then withdrawn. H2B-mCherry fluorescence was monitored using an IncuCyte live-cell imager (n=3). Dotted lines indicate conditions in which Dox was not administered . **D-E.** FACS-based assessment of Dox-mCherry label retention in GSC-0827 and GSC-464T cultures after 2, 5, and 9 days of chase. **F.** FACS analysis of Dox-mCherry GSC-0827 tumors after Dox withdrawal. Doxycycline (Dox) was administered in the drinking water either continuously or withdrawn for the indicated number of days prior to tumor harvest. It should be noted that most cells that we were not able to label the majority of cells due transgene silencing of the H2B-mCherry construct (which we consistently observe regardless of the LV-insert). **G.** FACS analysis of label retention using EdU incorporation in GSC-0827 tumors 1 and 5 days post EdU injection. **H.** As in **G** with additional replicates and time points.

However, the experiment and results should be considered in the context of the framework for GBM cell heterogeneity proposed by Lan et al (2017) for X-GBM tumors. From modeling of clone structure within GBM tumors, Lan and colleagues proposed a GBM hierarchy of slow-dividing stem-like cells giving rise to highly proliferative progenitors (via strict asymmetric division), which in turn give rise to non-dividing or dead progeny. If true, for our LR experiments, it is important to label both slow dividing stem-like cells as well as fast dividing progenitors. For H2B-mCherry labeling, we induced for 1 week prior to chase, which we estimate should be sufficient to label a significant portion of the slow dividing stem cells and all of the progenitors (not shown).

For both H2B-mCherry and EdU (Fig. 5), we observe continuous dilution of the label (in vitro: Fig. 5C, D, and E; in vivo Fig. 5F, G. and H). After 8-12 days only a small subset of LR+ cells persisted, accounting for less than 5% of the total tumor population by day 12. Note that due to transgene silencing of the vector, which is more prevalent in vivo (with this and other vectors), we could only achieve ∼20% labeling of the total tumor cell population after 1 week of induction of H2B-mCherry. Thus, although day 12 tumors had .97% labeled cells we assume that the actual number is closer to 5% total. However, the label retaining population still exhibited progressive label dilution, consistent with slower - but continuously dividing populations. This indicates that X- GBM Q states are transient and that nearly all tumor cells undergo division within 8 days and that those persisting in Q states are still dividing. This is consistent with scRNA-seq analysis of day 8 LR tumor cells (Fig. 1I), where although cells accumulate in Q states, cycling cells are still observed, albeit at lower frequencies. In general, these data suggest that that X-GBM tumors do not contain reservoirs of dormant Q cells and further support that notion that GBM Q states are more similar to primed or activated NSCs.

### Proliferative heterogeneity maybe sufficient to account for cell diversity in GBM tumors

If GBM tumors lack true reservoirs of dormant stem cells and do not recapitulate developmental programs such as adult neurogenesis, then their cellular heterogeneity must arise by other means. Perhaps tumor diversity could emerge not from fixed hierarchies but from proliferative heterogeneity itself? We, thereby, hypothesized that GBM tumors evolve into developmentally partitioned proliferative compartments (DPPCs) with complete cell cycles, transient Q states, and integration of Q, A, and differentiation networks, which collectively generate the spectrum of tumor cell states.

To test this hypothesis, we created a computational pipeline to assess proliferative heterogeneity in individual patient tumor samples (Figure 6A and B). The pipeline starts by restricting a scRNA-seq dataset down to the S-phase cells as defined by the ccAFv2 cell cycle classifier. Next, the DPPCs are identified by de novo clustering of the S-phase cells and subsequent differential expression analysis discovers marker genes that uniquely describe each DPPC. Overlap analysis of the marker genes from the DPPCs from all samples is used to determine the unique set of DPPCs across all the tumors in the study, and each tumors DPPCs are labeled such that DPPCs shared across tumors have the same label (e.g., DPPC1). This is extensible in that as new data is collected it can be easily integrated and its DPPCs labeled, and novel DPPCs added to the compendium. Next, we used deep learning methods to construct a classifier for each scRNA-seq dataset trained on the DPPCs labels and the expression of their marker genes. This classifier was then applied to the whole scRNA-seq dataset to provide DPPC labels beyond just the S phase cells. This DPPC classification approach does not require prior knowledge and thus should better account for patient-specific differences in tumor subpopulations. One important caveat is that, based on multiple pilot studies, at least 100 S-phase cells must be present in samples for the classifier to work. As a result, we initially focused on scRNA-seq data from 5 early passage GBM sphere cultures (7 days post isolation)(Couturier et al. 2020), which allowed sufficient numbers of cells.

**Figure 6.**
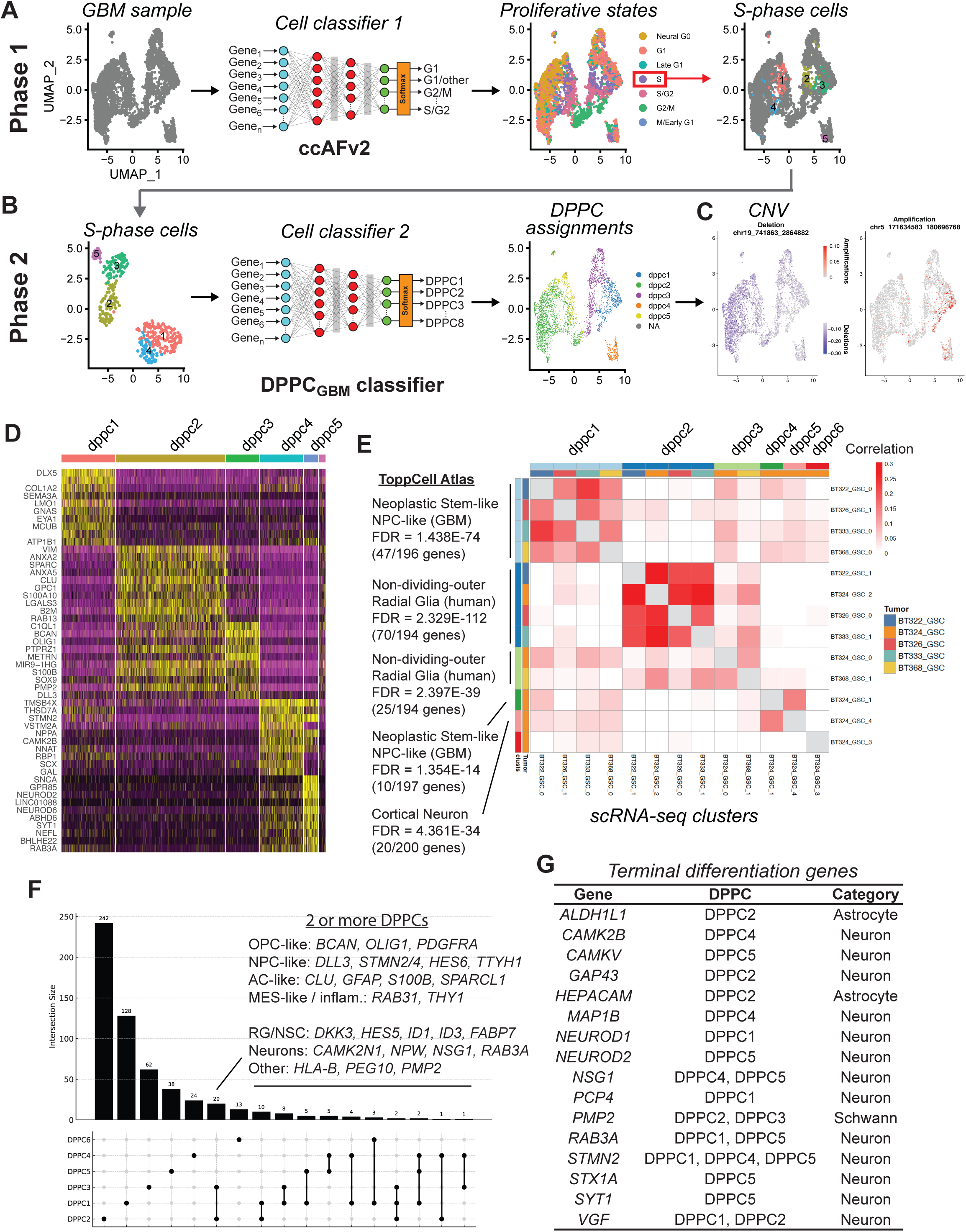
Overview and use of the DPPC classifier. **A & B.** Overview of DPPC classifier scheme for analyzing scRNA-seq datasets from GBM isolates. **C.** CNV analysis of BT324 scRNA-seq data (independent of DPPC assignments). **D.** Heat map of top 10 genes enriched in each DPPC for BT324 (from Fig. 1A, B, and C). **E.** Correlation of genes in DPPCs and their presence in de novo clusters from scRNA-seq data of GBM isolates shown, as well as ToppCell Atlas correlations for cell type for DPPCs. Supplemental Table S5 contains genes associated with DPPCs 1-6. Supplemental Table S6 contains overlap of genes associated with DPPCs 1-6 and scRNA-seq clusters shown. **F.** An upset plot showing overlap of genes within DPPCs. **G.** Representative terminal differentiation genes found within DPPCs.

The results for the BT324 GBM culture are shown in Figures 6A, B, and C. For this isolate, 5 DPPCs were identified, accounting for 90% of cells in the data set. DPPC-specific CNV alterations in Chr5 (DPPC1) and Chr19 (DPPC2) along with DPPC-specific gene expression patterns (Fig 6C, D, and E) supported the DPPC designations.

Comparing DPPCs across the 5 isolates revealed that most had 2 dominant DPPCs, and 3 DPPCs had common features between isolates (Fig. 6D and E) (Tables S5 and S6). These included DPPCs 1, 2, and 3 which had high proportions of gene overlapping with NPC and radial glial cells, cell type enrichment analysis using the ToppCell Atlas of human cell types (Fig. 6E). DPPCs also have overlapping expression of genes previously defined GBM cell states (e.g., Neftel et al subpopulations Ac-like, OPC-like, NPC-like, MES) (Fig. 6D). However, they are not necessarily defined by these markers which can be shared with more than one DPPC (Fig. 6F). In addition, DPPCs express genes expressed in terminally differentiated CNS cell types (Fig. 6G), including genes only predominantly expressed in post-division neurons (e.g. CAMKV, GAP43, RAB3A, SYT1 (Human Protein Atlas; www.proteinatlas.org). Thus, DPPCs populations of cells with admixtures of Q, A, and differentiation gene expression networks. While preliminary, the findings support the general notion that evolution of proliferative compartments with transient Q states and complete cell cycles may ultimately drive tumor cell diversity in GBM tumors.

## Discussion

Here, we computationally and functionally profiled Q populations in GBM xenograft and primary patient tumors using scRNA-seq data and in vivo phenotypic assays. Our results recast GBM Q as a transient, hybrid state embedded within proliferative compartments rather than a dormant stem-cell reservoir. From scRNA-seq and velocity mapping of functional defined Q cells found cell cycle egress points that flow toward OPC-like or RG-like programs. DNA label retention assays confirmed these termination points indeed harbor Q cells, but that even the longest Q-residing cells divide continuously. Transcriptional network analyses revealed that these Q populations co-activate canonical Q regulators of NSC (FOXO1, HES1, STAT3, SOX2/9) together with NSC A networks (ERG1/2, JUN, SOX8, etc.) that prime cell cycle re-entry. The same Q+A “hybrid” signature was evident in primary GBMs and included additional pro-proliferative TF networks (e.g., GLI2, HOXA/PBX1, LEF1, ONECUT2, ZNF217).

Motivated by these observations, we developed a patient-specific classifier that seeds on S-phase heterogeneity to define Developmentally Partitioned Proliferative Compartments (DPPCs) (Fig. 7A). Though preliminary, the successful identification of DPPCs suggests that tumors are composed of multiple proliferative compartments that have full cell cycles, transient Q states, and admixtures of Q, activation, and differentiation gene expression networks.

**Figure 7.**
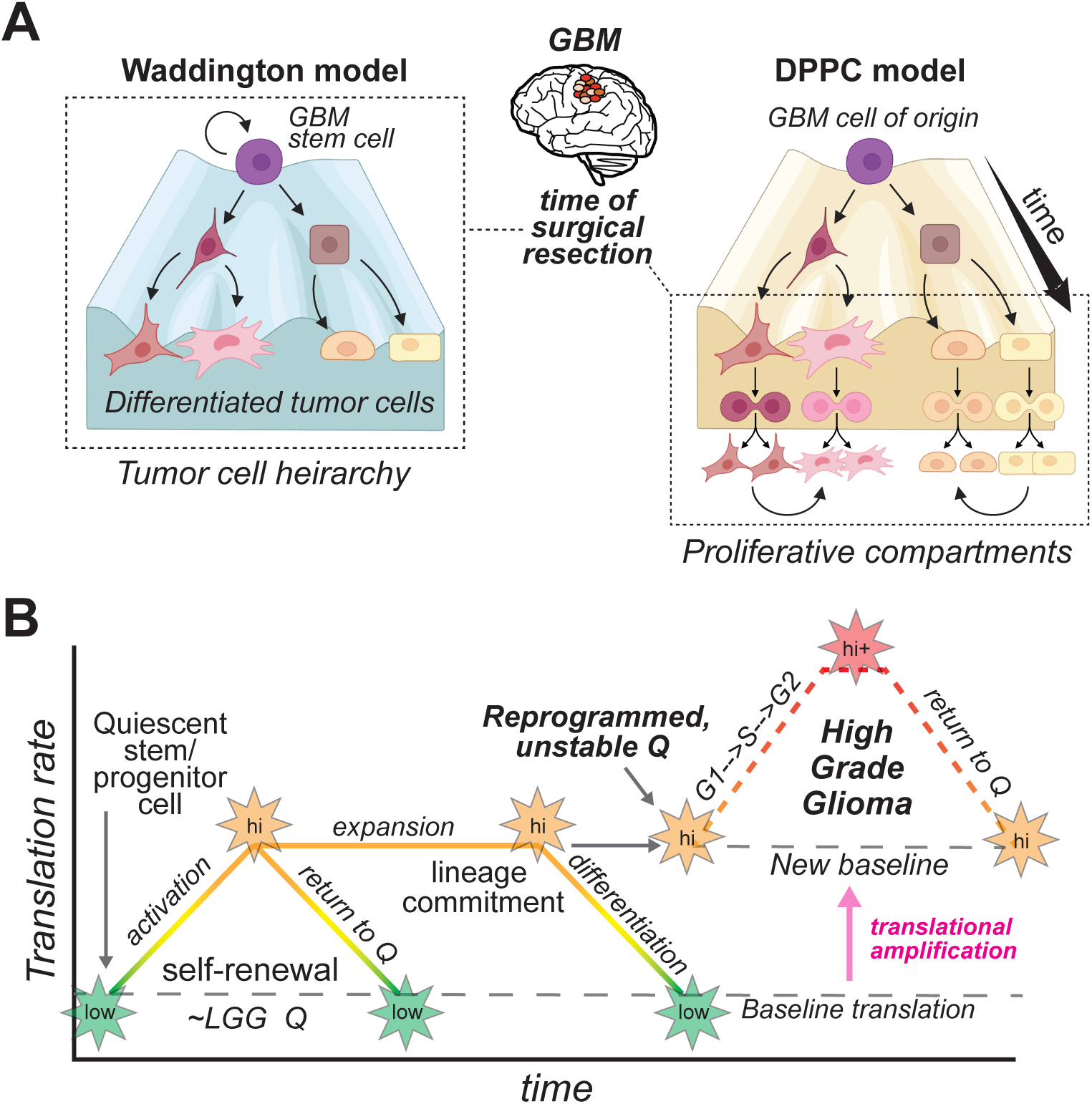
Overview of DPPC model and hypothesis that translational amplification drives Q instability. **A.** A model for DPPCs. **B.** A cartoon comparing translation rates in non-transformed NSCs, LGG and HGG.

This view diverges from prevailing notions that GBM tumors are hierarchically organized (Lathia et al. 2015). In canonical models, dormant CSCs initiate lineages analogous to neurogenesis, progressing from Q stem cells to activated progenitors, amplifying progenitors, and ultimately differentiated progeny. More recent quantitative models are less strict but maintain these general categories. Lan et al. (2017) used DNA barcode-based lineage tracing of clonal outgrowth human xenograft tumors to propose a quantitative model of slow-cycling GBM stem-like cells that undergo invariant asymmetric divisions to feed a rapidly cycling progenitor pool with limited growth potential. However, this model does not account for several key biological variables, including lineage plasticity, “equipotency”, and niche-conditioning which could give rise similar data without a hierarchy.

For example, from our scRNA-seq data, GSCs adopt states not observed in vitro cultures, that are associated with slower dividing Q like states (Mihalas et al. 2025). For the GSC-0827 isolate, a GFAP+ population only emerges in vivo, and although primarily confined to a Q cluster (cluster 8 in reference data (Fig. 1A)), GFAP+ cells nonetheless have a full cell cycle but with lower frequencies of S/G2/M cells, suggestive of slower G1-S entry. It seems likely that post-engraftment transitions governed by niche-driven BMP/TGFb-SMAD signaling give rise to GFAP+ cells (Sachdeva et al. 2019; Anido et al. 2010)). [In vitro this states can be reversibly induced with serum (not shown)(Carén et al. 2015).] If true, lineage plasticity coupled with microenvironmental signaling could give rise to similar clonal dynamics observed by Lan and colleagues. Thereby, transient Q and hybrid Q-A states likely better fit with a model whereby transitions into and out of longer Q states is driven by niche-dependent and also oncogenic rewiring of key cell intrinsic factors.

One such cell intrinsic factor is translation rate. Protein translation is limiting for exit from Q -- as artificially increasing translation is sufficient to trigger Q egress/cell cycle entry even in the absence of growth factors (Smith et al. 1990) by promoting translation of key cell cycle entry genes, including cyclin D1 (Rosenwald et al. 1995). We recently found that prospective Q states in primary GBM tumors have amplified translation rates that are approximately equivalent to the highest rates observed in low grade gliomas and also NSCs (Mihalas et al. 2025) (Fig. 7B). This new supraphysiological basal-line for translation is dependent on the protein acetyl transferase KAT5 (Mihalas et al. 2025). Accompanying increased translation, KAT5 coordinates transcription of NSC A TF networks, including AP-1, E2F, and MYC (Mihalas et al. 2025). Further, serially decreasing KAT5 activity results in proportional decreases protein translation rates and increases in steady-state Q populations (Mihalas et al. 2025).

For GBM Q cells, aside from KAT5-driven transcriptional effects, global translation amplification may ultimately drive Q instability (Fig. 7B). For it could bias the proteome to favor the activity of otherwise labile oncoproteins proteins and signaling of A TF networks (e.g., AP-1, E2F, MYC, Cyclin D (short ∼<1hr half-lives)) found in hybrid Q-A states. By contrast, Q networks are likely specificized in part by longer-lived TFs that appear in Q networks, such as SOX2 and SOX9 (∼7-8hr half-lives (Strebinger et al. 2019; Hong et al. 2016)), similar to other developmental TFs (e.g. (Gurdon et al. 2020)). Thus, changes in global translation rates could ultimately drive “conflicts” between developmental circuits controlling indolent Q program and A networks driving aggressive growth. The latter is normally kept in check by tumor suppressive pathways (e.g., Rb-axis) and Q niche-dependent feedback control, which are compromised in tumors.

In sum, our results suggest that GBM Q is a transient, hybrid state likely destabilized by translational amplification, rather than a reservoir of dormant stem cells. We propose that proliferative heterogeneity organized into dynamic compartments, not fixed CSC hierarchies, underlies GBM diversity and therapeutic resistance.

## Materials and Methods

### Ethical Statement

This research was approved by the Fred Hutch Institutional Animal Care and Use Committee (Protocol # 202100013) and complies with all required ethical regulations.

### Cell Culture

Patient tumor-derived GSCs were provided by Drs. Jeongwu Lee (Cleveland Clinic), Do-Hyun Nam (Samsung Medical Center, Seoul, Korea), and Steven M. Pollard (University of Edinburgh). Isolates were cultured in NeuroCult NS-A basal medium (StemCell Technologies) supplemented with B27 (Thermo Fisher Scientific), N2 (homemade 2x stock in Advanced DMEM/F-12 (Thermo Fisher Scientific)), EGF and FGF-2 (20 ng/ml) (PeproTech), glutamax (Thermo Fisher Scientific), and antibiotic-antimycotic (Thermo Fisher Scientific). Cells were cultured on laminin (Trevigen or in-house-purified)-coated polystyrene plates and passaged as previously described^97^, using Accutase (EMD Millipore) to detach cells.

### GSC Tumors

NSG mice (Jackson Labs #005557) used in this study were kept in a 12h light/dark cycle at 72°F and 40-60% relative humidity with food and water ad libitum, in the Fred Hutchinson Cancer Center (FHCC) vivarium. All animal experimental procedures were performed with approval of the FHCC Institutional Animal Care and Use Committee (Protocol # 202100013). All procedures followed guidelines outlined in the National Research Council Guide for the Care and Use of Laboratory Animals. 100,000 GSCs were orthotopically xenografted into a single frontal cerebral hemisphere. GSCs were injected using stereo-tactic coordinates: 2 mm lateral from Bregma and 3.5 mm depth and grown for 3-12 weeks according to our previously published protocols^73,98,99^. We relied on MRI and clinical/neurological symptoms to evaluate the tumor burden. Once the tumor was visible by MRI, mice were monitored 3 times a week and then daily if neurological symptoms were observed. Animals that exhibited the following criteria for maximal tumor burden were euthanized and considered at endpoint: impaired mobility, inability to reach food and water, hunched posture, labored breathing and/or cyanosis (bluish ears, feet or mucous membranes), abnormal response to stimuli (slow to move/does not move or reacts excessively), body conditioning, and skin ulceration and/or necrosis, in the event of extracranial tumor growth. Maximal tumor burden was not exceeded in these studies. *<u>Doxycycline dosage of mice</u>:* After tumors reached 2mm^3^ by MRI, NSG mice were dosed with Doxycycline (2 mg/ml) in the drinking water (supplemented with 5% sucrose) for 1 week and then grouped experimental cohorts (e.g., continued Dox and Dox removal). *<u>EdU pulsing of mice:</u>* Between 24h and 8 days prior to tumor harvesting, mice were intra-peritoneally injected with EdU (100 mg/kg) and tumors were harvested between 1 and 8 days later. Cells were analyzed using the Click-iT™ EdU Flow Cytometry Assay Kit (Thermo Fisher Scientific) according to the manufacturer’s protocol. *<u>MRI:</u>* volume of interest was manually contoured using T2-weighted brain MRI scans (Bruker 1T and 7T scanner) for all animals using Horos Dicom Viewer software (horosproject.org).

### p27 and LR Reporters

The p27 reporter was constructed after (Oki et al., 2014), using a p27 allele that harbors two amino acid substitutions (F62A and F64A) that block binding to Cyclin/CDK complexes but do not interfere with its cell cycle-dependent proteolysis. This p27K^-^ allele was fused to mVenus to create p27K^-^-mVenus and cloned via Gibson assembly (NEB) into a modified pGIPz lentiviral expression vector (Mihalas et al. 2025). P27-mVenus reporter cells were sorted for the presence of mVenus on an FACSAria II (BD) and normal growth was verified post-sorting.

To introduce the H2B-mCherry label retention reporter into cells, the G0LR plasmid (where H2B-mCherry ORF was cloned from Addgene vector #20972) into was cut with EcoRV to generate double stranded DNA fragments with homology to the SHS231 safe harbor locus (Pellenz et al. 2019). RNP complexes were formed with a chemically synthesized sgRNA (Synthego) targeting the SHS231 safe harbor locus and purified sNLS-SpCas9-sNLS (Aldevron) (Cas9:sgRNA ratio of 1:2). dsDNA and RNP complexes were electroporated into cells using the Amaxa 96-well Shuttle System (Lonza) and program EN-138 (as described in (Hoellerbauer et al. 2020)). For in vivo experiments, cells were infected with lentivirus generated from the G0LR plasmid. After G0LR integration into cells, cells were selected with G418 for 5 days. Cells went through 2 rounds of sorting to exclude false positive and false negative reporter events: cells were grown without Doxycyclin and mCherry negative cells were sorted for and expanded in culture, cells were then treated with Doxycyclin for 2 days and mCherry+ cells were sorted for and expanded in culture for 1+ weeks prior to experiments. To create clones, single G0LR cells were plated in 96 well plates with an additional 2,000 parental cells. After growth for 7 days, G0LR cells were selected with G418 for 5 days followed by expansion to generate cell numbers for experiments.

### Lentiviral Production

For virus production, lentiCRISPR v2 plasmids were transfected using polyethylenimine (Polysciences) into 293T cells along with psPAX and pMD2.G packaging plasmids (Addgene) to produce lentivirus. For the whole-genome CRISPR-Cas9 libraries, 25x150mm plates of 293T cells were seeded at ∼15 million cells per plate. Fresh media was added 24 hours later and viral supernatant harvested 24 and 48 hours after that.

For screening, virus was concentrated 1000x following ultracentrifugation at 6800*xg* for 20 hours. For validation, lentivirus was used unconcentrated at an MOI<1.

### scRNA-seq analysis

Single cell RNA-sequencing was performed using 10x Genomics’ reagents, instruments, and protocols. Single cell RNA-Seq libraries were prepared using Chromium Single Cell 3ʹ Reagent Kit. CellRanger (Zheng et al. 2017) ( v5.0 from 10× Genomics) was used to align, quantify, and provide basic quality control metrics for the scRNA-seq data.

Souporcell (Heaton et al. 2020) was used to deconvolute scRNA-seq data for each GSC cell line. Using Seurat (Butler et al. 2018) (version 4), the scRNA-seq data was normalized using the SCTransform pipeline and were merged the tumor replicates to build an integrated reference. FindTransferAnchors and MapQuery from Seurat, was used to map the query tumors to the integrated reference. FindAllMarkers was used to find differentially expressed genes for each cluster of each tumor. AddModuleScore from Seurat to calculate the average expression levels of different gene lists of interest for each tumor type. ggplot2 was visualize to make bar plots to visualize the number of genes and cells in each cluster. ccSeurat (Butler et al. 2018) and ccAF (O’Connor et al. 2021) were used to score cell cycle states for each cell. scVelo^111^ was used to perform velocity analysis. The ToppCell Atlas (Jin et al. 2021) was used to perform gene set enrichment analysis on each of the differentially expressed gene list from each cluster.

For scRNA-seq analysis, >1000 cells was used for each scRNA-seq analysis with a per cell sequence read depth of >20,000. Standard Seurat filters were applied requiring that the cells had to have at least 200 features per cell, and transcripts need to be expressed in at least three cells. Each sample was further filtered by requiring the number of UMIs per cell to fall within a range of UMIs and the mitochondrial percentage of genes expressed per cell to fall within a range of percents. The cells from each sample were then normalized using SCTransform, principal components were calculated, and a UMAP was generated. To identify UMAP gene cluster markers, a Wilcoxon rank sum test and Bonferroni correction were used. To compare cluster-specific gene expression (i.e., differential expression analysis), negative binomial tests (DESeq2) and Benjamini-Hochberg False Discovery Rate (FDR≤ 0.05) corrections were used. The bioinformatics analysis was processed at least twice, and results were compared to ensure the reproducibility of our findings.

### Discovering TF regulators from scRNA-seq cluster marker genes

The input for the TF regulator discovery pipeline are the marker genes for a scRNA-seq cluster (average log2 fold-change ≥ 1 and Bonferroni adjusted p-value ≤ 0.05). First, the enrichment of binding sites from the TFBSDB (Plaisier et al. 2016) was used to establish mechanistic evidence of TF regulation. A hypergeometric p-value was computed for the overlap between each TF regulator in TFBSDB versus the marker genes from the cluster using a hypergeometric p-value. Significant enrichment of a TF binding sites was observed with a Benjamini-Hochberg corrected p-value (q-value) ≤ 0.05 and percent of targets greater than zero. Then, the TF family expansion method used in Plaisier et al., 2016 was used to extend the likely TF candidate regulators based on similarity of binding motifs among family members from the TFClass database [REF: 29087517]. Then the expression of the TF was correlated (Pearson’s) with the expression of the marker genes with binding sites summarized as an eigengene (first principal component corrected for sign) (Langfelder and Horvath 2008; Plaisier et al. 2016). Associations between TFs and their marker genes were considered significant if the Benjamini-Hochberg corrected p-value (q-value) was ≤ 0.05 and the correlation coefficient was greater than zero.

### Discovering DPPCs

First, we applied the ccAFv2 cell cycle classifier (O’Connor et al. 2025) to the BT324 GSC scRNA-seq dataset (Couturier et al. 2020). Then we restricted our analysis to the S phase cells based on the ccAFv2 predictions to control out the cell cycle signal and allow us to discover the distinct expression patterns underlying our cycling cells. We then used de novo clustering in Seurat to discover the unique sets of cells with distinct expression patterns. There clusters of S phase cells with distinct expression signatures are the DPPCs. Next, we used the S phase cells and DPPC identities to train an artificial neural network classifier with the same structure as ccAFv2 (O’Connor et al. 2025), except for the number of input and output nodes which were set to the number of significant marker genes for the DPPCs and number of DPPCs, respectively. The classifier was then applied to all the cells in BT324 GSC scRNA-seq dataset to predict the DPPC for every single cell. DPPCs from the six GSC lines from Couturier et al., 2020 were applied with the same pipeline and comparison of the marker genes were made and significance determined by hypergeometric overlap analysis.

#### Data Availability

The scRNA-seq data files are available on the GEO database at GSE198524 and GSCXXXX.

#### Code Availability

The code used to process and analyze the data is available at https://github.com/sonali bioc/GSC_scRNASeq; https://github.com/plaisier-lab/.

## Supporting information

Suppl Inventory and Figures

## Acknowledgments

We thank members of the Paddison, Plaisier, and Patel labs for helpful discussions, Dr. Atsushi Miyawaki for providing reagents, and Annique Lennon for administrative support. This work was supported by the following grants: Fred Hutch pilot award (A.P., P.P.); NCI/NIH (R01CA295090) (P.P.); NINDS/NIH (R01NS119650) (A.P., C.P., P.P.); Burroughs Wellcome Career Award for Medical Scientists (A.P.); and a grant from the Kuni Foundation (A.P.). Preclinical Imaging Core, Fred Hutchinson Cancer Center support: P30 CA015704 (RRID:SCR_022616); 3T/7T MRI SIG: NIH S10 OD26919.

## Author contributions

Project conception and design was carried out by P.P., C.P., A.P., A.M., and K.M.. Experiments and data analysis were performed by A.M., and K.M. Single cell bioinformatical data analysis and statistics were performed by S.A., S.O., and C.P. with input from A.P., C.P., and P.P.; P.P. and C.P. wrote the manuscript with input from other authors.

