## Supplementary material for "Characterization of quiescent subpopulations and proliferative compartments in glioblastoma": Suppl Inventory and Figures

**Supplemental Figures**

**Supplemental Figure S1**: Data in support of Figure 1 and Figure S2, cut offs used for scRNA-seq data for p27hi GSC xenograft tumors.

**Supplemental Figure S2**: Analysis of Q states in GSC-464T X-GBM tumors.

**Supplemental Figure S3**: mRNA velocity analysis of total and p27hi cell populations in GSC-0827 tumors, associated with Figure 1.

**Supplemental Figure S4**: Expression analysis of CCND1 and CCND2 in scRNA-seq data from GSC xenograft tumors by cluster, associated with Figure 3.

**Supplemental Tables**

**Supplemental Table S1**: The eigengene for TF targets used identify TFs with expression that correlates targets (associated with Figure 2A).

**Supplemental Table S2**: Cancer hallmark GO terms with semantic similarity scores of ≥ 0.80 for TFs from Table S1 (associated with Figure 2A).

**Supplemental Table S3**: TFRegMap analysis of adult mouse Q and A NSCs from Llorens-Bodadilla et al. (2015) (GSE67833) (associated with Figure 2B).

**Supplemental Table S4**: TFRegMap analysis of ccAF_2 defined Q populations from scRNA-seq primary GBM tumors (associated with Figure 4B).

**Supplemental Table S5**: List of genes associated with DPPCs 1-6 (associated with Figure 6).

**Supplemental Table S6**: Overlapping genes associated with DPPCs 1-6 (associated with Figure 6).

**Supplemental Figure Legends**

**Supplemental Figure S1:** Cutoff filters used for percentage of mitochondrial genes and total number of RNA molecules (or Unique Molecular Identifiers, UMIs) for p27hi scRNA-seq data sets from Figure 1 and Supplemental Figure S2.

**Supplemental Figure S2:** Analysis of Q states in GSC-464T X-GBM tumors.
**A-B.** scRNA-seq UMAPs for GSC-464T tumor reference and p27hi tumor cells. Overview of experiment, filter cutoffs, and QC analysis are available (Mihalas et al. 2025).
**C-D.** Relative proportions of mapped single cells appearing in clusters from **A** and **B.**
**E.** mRNA Velocity analysis (scVelo) of p27hi cells reveals trajectories toward Q states after mitotic exit.
**F.** ccAF classification of p27hi cells.
QC cut offs for scRNA-seq analysis is available in Supplemental Figure S1.

**Supplemental Figure S3:** mRNA velocity analysis of total and p27hi cell populations in GSC-0827 tumors.
**A-B**, RNA velocity analysis of scRNA-seq data for **Figure 1**.
**C-E**, Key cell cycle transcription factors with significant mRNA velocity differences in GSC-0827 scRNA data.
**C**, This column of graphs shows unspliced versus spliced ratio of candidate genes with significant changes in velocity between clusters, where each data point is a cell colored by cluster from **A**. Contoured lines are derived from dynamical modeling of mRNA velocity from scVelo (https://scvelo.readthedocs.io/) (Bergen et al., 2020).
**D,** Velocity values for each gene mapped onto UMAP projection of GSC-0827 tumor reference. High velocity values for a particular gene (i.e., higher predicted unspliced/spliced mRNA ratios) tend to occur before expression peaks, followed by lower velocity and transcriptional down regulation.
**E**, Gene expression values for each gene mapped onto UMAP projection of GSC-0827 tumor reference.

**Supplemental Figure S4:**Expression analysis of *CCND1* and *CCND2* in scRNA-seq data from GSC xenograft tumors by cluster, associated with Figure 3.

### Figure S1

**A** **X-GSC-0827**
**p27$^{hi}$ tumor cells**

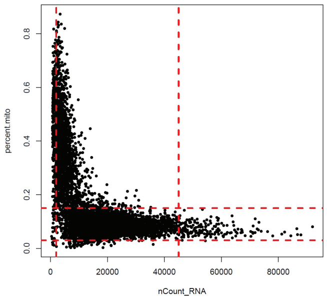


**B** **X-GSC-464T**
**p27$^{hi}$ tumor cells**

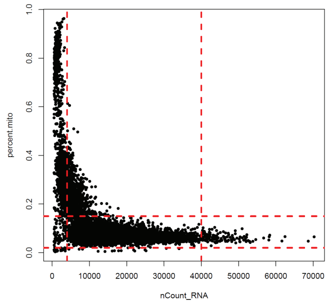


<u>Filters applied</u>:
0.03 < percent.mito < 0.15
2000 < nCount_RNA < 45000

### Figure S2

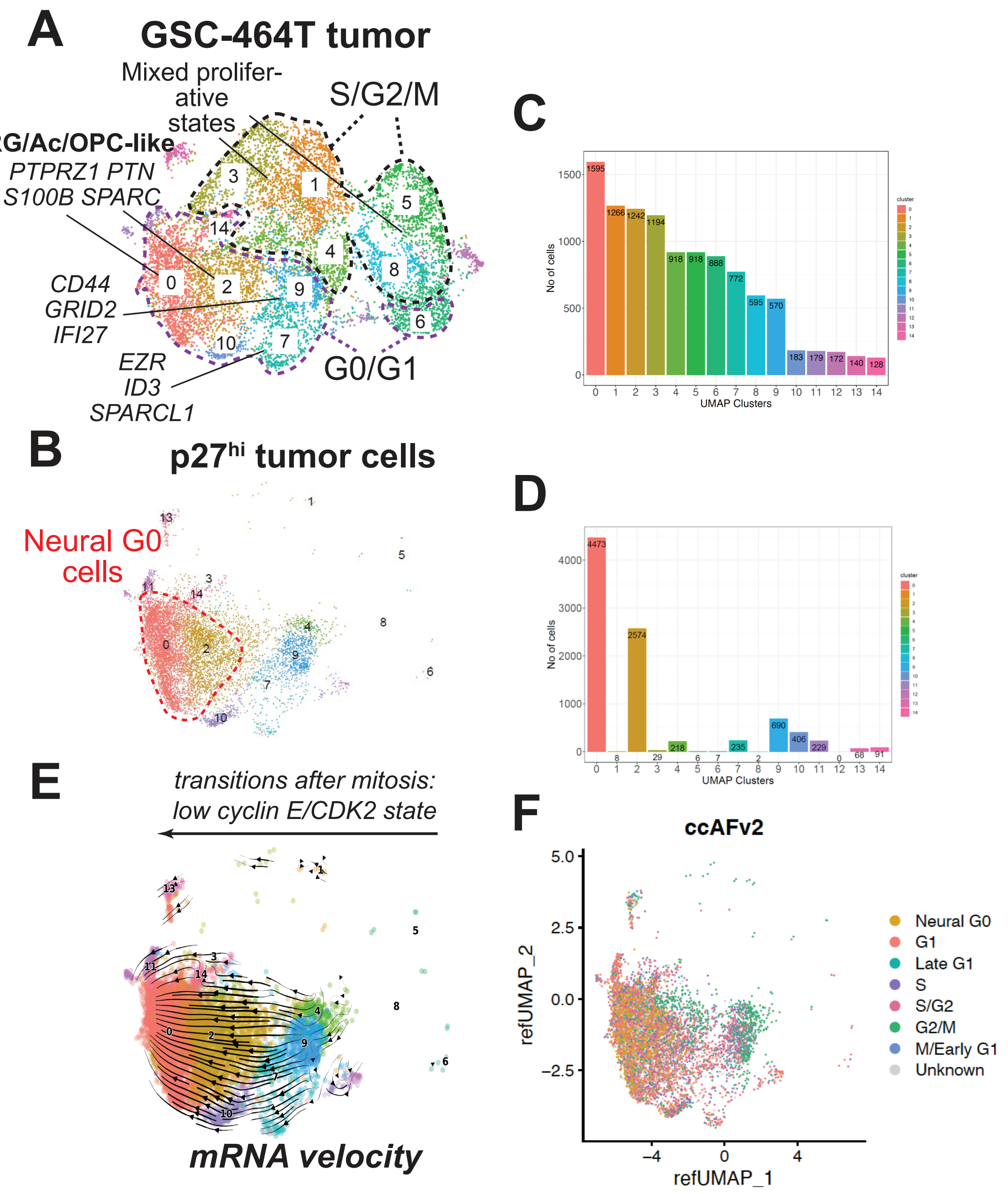

A
GSC-464T tumor
Mixed prolifer-
ative
states
S/G2/M
RG/Ac/OPC-like
PTPRZ1 PTN
S100B SPARC
CD44
GRID2
IFI27
EZR
ID3
SPARCL1
G0/G1
B
p27hi tumor cells
Neural G0
cells
C
No of cells
UMAP Clusters
cluster
D
No of cells
UMAP Clusters
cluster
E
transitions after mitosis:
low cyclin E/CDK2 state
mRNA velocity
F
ccAFv2
refUMAP_2
refUMAP_1
Neural G0
G1
Late G1
S
S/G2
G2/M
M/Early G1
Unknown

### Figure S3

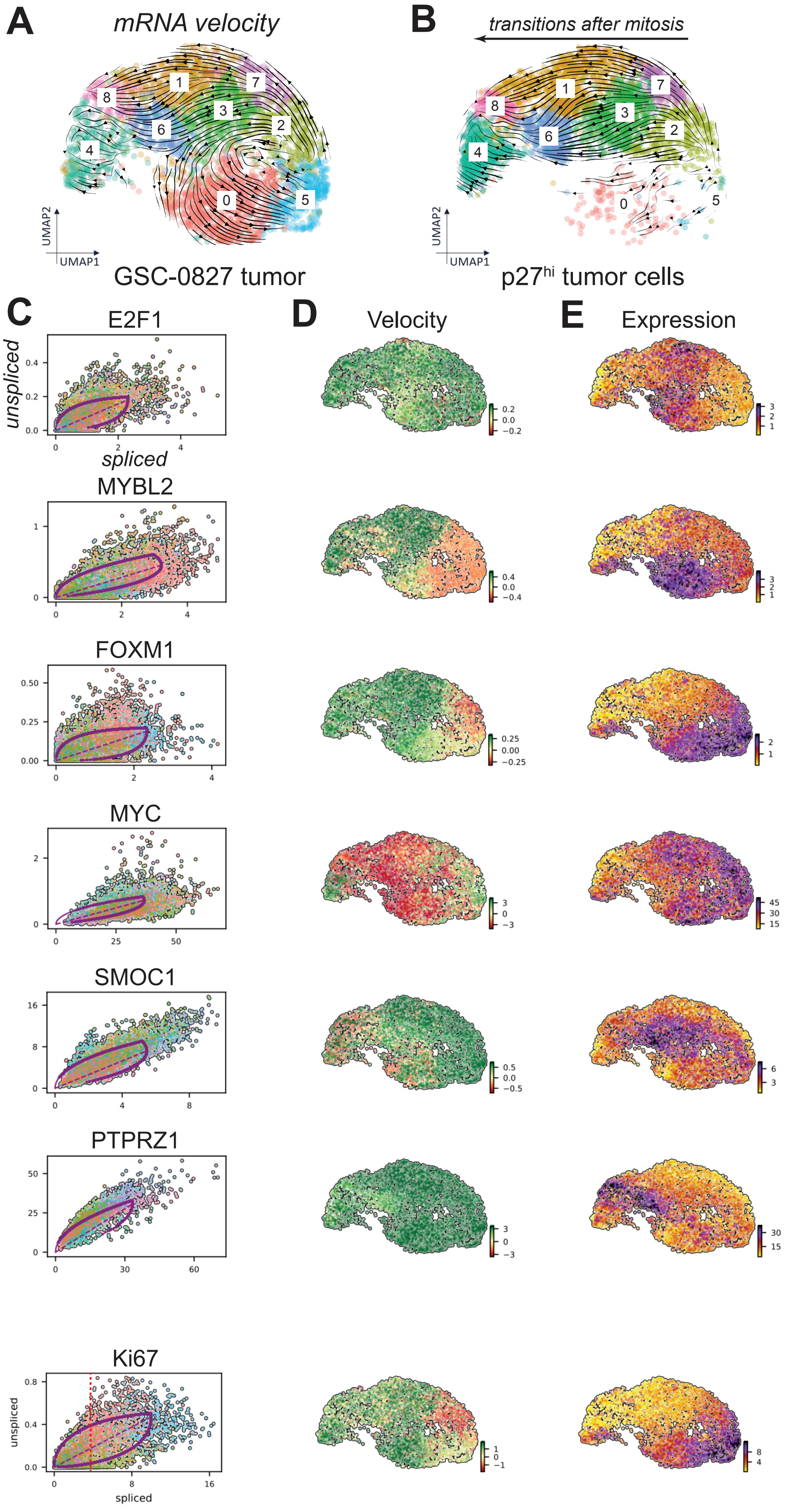
A
mRNA velocity
B
transitions after mitosis
UMAP2
UMAP1
GSC-0827 tumor
p27hi tumor cells
C
D
Velocity
E
Expression
E2F1
unspliced
spliced
MYBL2
FOXM1
MYC
SMOC1
PTPRZ1
Ki67

### Figure S4

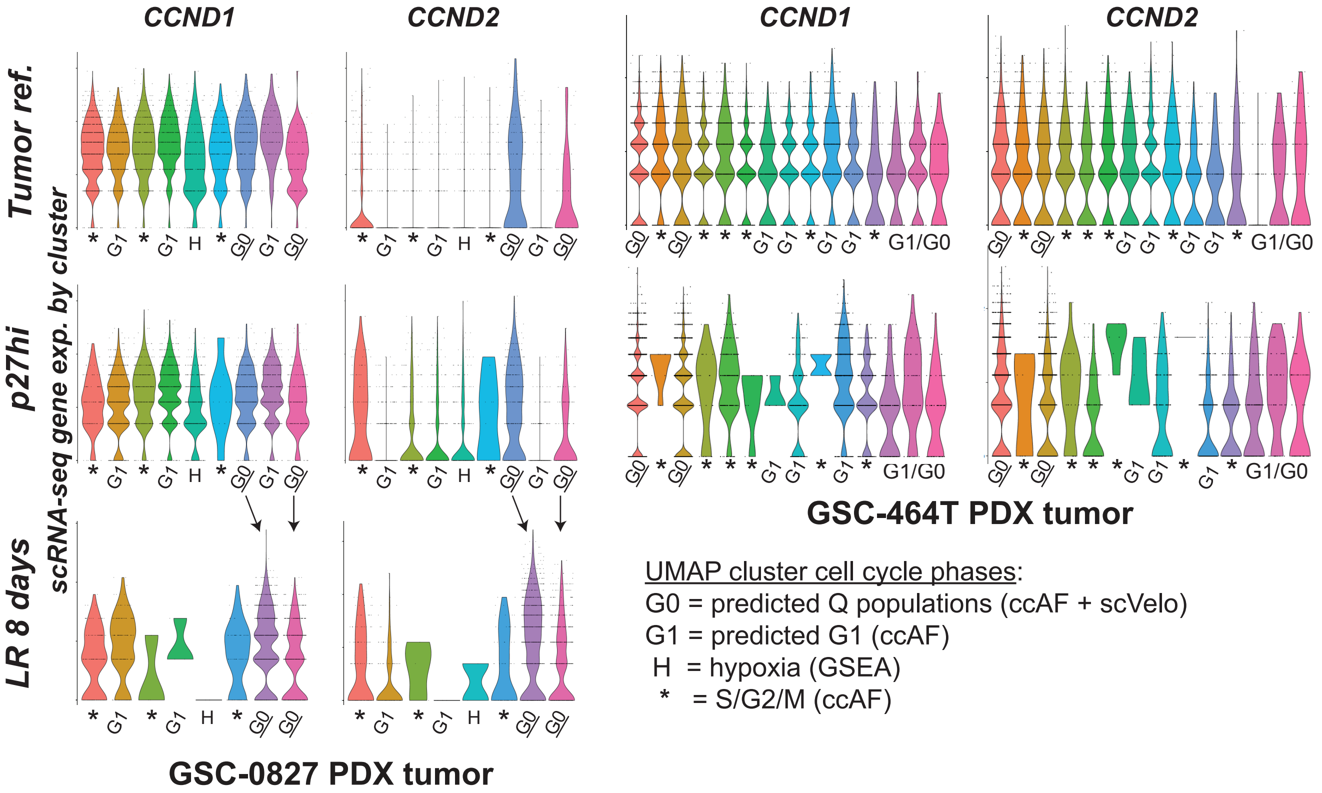

CCND1
CCND2
CCND1
CCND2
Tumor ref.
p27hi
LR 8 days
scRNA-seq gene exp. by cluster
* G1 * G1 H * G0 G1 G0
G0 * G0 * * * G1 G1 * G1 G1 * G1/G0
G0 * G0 * * * G1 G1 * G1 G1 * G1/G0
G0 * G0 * * * G1 G1 * G1 * G1/G0
G0 * G0 * * * G1 G1 * G1 * G1/G0
* G1 * G1 H * G0 G0
GSC-464T PDX tumor
GSC-0827 PDX tumor
UMAP cluster cell cycle phases:
G0 = predicted Q populations (ccAF + scVelo)
G1 = predicted G1 (ccAF)
H = hypoxia (GSEA)
* = S/G2/M (ccAF)
